# Atazanavir is a competitive inhibitor of SARS-CoV-2 M^pro^, impairing variants replication *in vitro* and *in vivo*

**DOI:** 10.1101/2021.11.24.469775

**Authors:** Otávio Augusto Chaves, Carolina Q. Sacramento, André C. Ferreira, Mayara Mattos, Natalia Fintelman-Rodrigues, Jairo R. Temerozo, Douglas Pereira Pinto, Gabriel P. E. da Silveira, Laís Bastos da Fonseca, Heliana Martins Pereira, Aluana Santana Carlos, Joana da Costa Pinto d’Ávila, João P.B. Viola, Robson Q. Monteiro, Leonardo Vazquez, Patrícia T. Bozza, Hugo Caire Castro-Faria-Neto, Thiago Moreno L. Souza

## Abstract

Atazanavir (ATV) has already been considered as a potential repurposing drug to 2019 coronavirus disease (COVID-19), however, there are controversial reports on its mechanism of action and effectiveness as anti-severe acute respiratory syndrome coronavirus 2 (SARS-CoV-2). Through the pre-clinical chain of experiments: enzymatic, molecular docking, cell-based, and *in vivo* assays, it is demonstrated here that both SARS-CoV-2 B.1 lineage and variant of concern gamma are susceptible to this antiretroviral. Enzymatic assays and molecular docking calculations showed that SARS-CoV-2 main protease (M^pro^) was inhibited by ATV, with Morrison’s inhibitory constant (Ki) 1.5-fold higher than boceprevir (GC376, a positive control). ATV was a competitive inhibition, increasing the M^pro^’s Michaelis-Menten (K_m_) more than 6-fold. Cell-based assays indicated that SARS-CoV-2 gamma is more susceptible to ATV than its predecessor strain B.1. Using oral administration of ATV in mice to reach plasmatic exposure similar to humans, transgenic mice expression in human angiotensin converting enzyme 2 (K18-hACE2) were partially protected against lethal challenge with SARS-CoV-2 gamma. Moreover, less cell death and inflammation were observed in the lung from infected and treated mice. Our studies may contribute to a better comprehension of the M^pro^/ATV interaction, which could pave the way to the development of specific inhibitors of this viral protease.

## 1. Introduction

The 2019 coronavirus disease (COVID-19) was firstly reported in Wuhan (China) and its etiological agent, severe acute respiratory syndrome coronavirus 2 (SARS-CoV-2), spread globally into the second pandemic of the 21^st^ century, after influenza A(H1N1) pandemic in 2009. SARS-CoV-2 continuously circulate, even among individuals with preexisting immunity, and caused about 258 million confirmed cases along with 5.16 million deaths worldwide [1-5]. SARS-CoV-2 variants of concern (VoC) might escape the humoral immune response to natural infection or vaccination [4,5], which reinforces the necessity of specific antiviral treatments. After almost a two-year effort in the repurposing of clinically approved drugs, limited benefit for COVID-19 patients have been demonstrated [5]. Thus, it is necessary to improve the pre-clinical characterization of repurposed drugs to rationalize further clinical studies, in terms of posology and susceptibility of VoC, and learn from their interactions with the viral target enzyme.

We have previously described that atazanavir (ATV), a clinically approved human immunodeficiency virus (HIV) protease inhibitor [6], is endowed with anti-SARS-CoV-2 activity [7]. Although we described early in the pandemic outbreak the ATV’s activity against SARS-CoV-2, by *in silico* and cell-based assays (in Vero-E6 and A549 cell lines), proper characterization of its enzymatic target, pharmacology on type II pneumocytes (the most affect cell in severe COVID-19) and antiviral activity in infected mice have not been described. Nevertheless, numerous clinical trials were initiated for out- and in-patients with COVID-19 to test ATV combined or not with other commercial drugs, such as ritonavir or dexamethasone [7-9], however no definitive response emerged from these clinical trials yet.

Studies reconfirmed, by bioinformatics and basic enzymatic inhibition curves, that ATV targets SARS-CoV-2 major protease (M^pro^), an enzyme responsible for the cleavage of eleven sites of the viral polyprotein, a key-step in virus life cycle [10,11]. The development of PAXLOVID™ from Pfizer and its clinical efficacy of reduce hospitalization by 80% reconfirms that M^pro^ is a very important druggable target [12]. Thus, other drugs that target this enzyme deserve a more detailed characterization, to provide insight on pharmacophoric regions for next generation of anti-COVID-19 antivirals.

Anti-M^pro^ from ATV has been disputed as controversial depending on assay conditions [13]. This apparent paradox reinforces that detailed mechanism of action and robust series of pre-clinical experiments should be conducted to shed light on the ATV inhibitory mechanism and the susceptibility of contemporaneous SARS-CoV-2 strains to this drug. Here, we address additional explanations to controversial effects of ATV on M^pro^, characterizing it as a competitive inhibitor of this viral enzyme that requires a catalytic water to be effective. ATV possesses anti-SARS-CoV-2 activity against B.1 and gamma strains on Calu-3 cells, a model of type II pneumocytes. ATV reached the plasma and lungs of treated Swiss-Webster mice and protected SARS-CoV-2-infected transgenic mice expression in human angiotensin converting enzyme 2 (K18-hACE2) from mortality. Moreover, ATV reduced virus-induced inflammation and cell death in bronchoalveolar lavage (BAL), and lung damage in infected and treated animals. This study compiles pre-clinical results that may allow further rationalization of clinical trials against COVID-19.

## 2. Materials and Methods

### 2.1. General materials

Carboxymethyl cellulose (CMC), *tris*(4-(dimethylamino)phenyl)methylium chloride (crystal violet), formaldehyde, hematoxylin, eosin, phosphate buffer solution (PBS), ethanol, dimethyl sulfoxide (DMSO), ketamine, xylazine, trisodium citrate, calcium chloride (CaCl_2_), and remdesivir (RDV) were purchased from Sigma-Aldrich/Merck (St. Louis, MO, USA). Atazanavir sulfate (ATV) was kindly donated from *Instituto de Tecnologia de Farmacos*, Farmanguinhos, Rio de Janeiro, Brazil.

### 2.2. Cells and virus

African green monkey kidney (Vero, subtype E6) and human lung epithelial (Calu-3) cells were cultured in high-glucose Dulbecco’s modified Eagle medium (DMEM - HyClone, Logan, Utah) supplemented with 100 U/mL penicillin, 100 μg/mL streptomycin (P/S - Thermo Fisher Scientific®, Massachusetts, USA), and 10% fetal bovine serum (FBS - HyClone, Logan, Utah). The cells were incubated at 310K in 5 % of carbon dioxide (CO_2_).

The SARS-CoV-2 B.1 lineage (GenBank #MT710714) and gamma variant (also known as P1 or B.1.1.28 lineage; #EPI_ISL_1060902) were isolated on Vero E6 cells from nasopharyngeal swabs of confirmed cases. All procedures related to virus culture were handled at biosafety level 3 (BSL3) multiuser facility at *Fundação Oswaldo Cruz* (Fiocruz), Rio de Janeiro, Brazil, according to World Health Organization (WHO) guidelines [14].

### 2.3. Enzymatic assays

The ATV capacity to inhibit enzymatic velocity of PL^pro^ and M^pro^ from SARS-CoV-2 was determined by the commercial kit provided by BPS Bioscience® company (catalog number: #79995-1 and #79955-1, respectively) following the procedure and recommendations from literature and manufacturer [15-17]. Basically, 100 nM PL^pro^ was incubated in 50 mM HEPES pH 7.4, 0.01% Triton X-100 (v/v), 0.1 mg/mL bovine serum albumin (BSA), 2 mM dithiothreitol (DTT) with 25 µM of its substrate (modified peptide Z-RLRGG-AMC with CAS number 167698-69-3) in the presence various concentrations (0-10 μM of ATV or GRL0617 (positive control) for 45-60 minutes. On the other hand, 88.8 nM M^pro^ was incubated overnight in reaction buffer (20 mM Tris pH 7.3, 100 mM NaCl, 1 mM EDTA, 1 mM DTT, and 1 μM BSA) containing 25 μM of substrate (modified peptide DABCYL-KTSAVLQSGFRKME-EDANS with CAS number 730985-86-1) and ATV or boceprevir (GC376, positive control), at concentrations ranging from 0 to 10 μM. Fluorescence signal was measured at an emission wavelength of 460 nm with excitation at 360 nm in a GloMax® (Promega) plate reader. Morrison’s inhibitory constant (K_i_) was calculated by non-linear regression using GraphPad Prism 9. The Michaelis-Menten plot was conducted for 88.8 nM M^pro^ incubated overnight in assay buffer with substrate concentrations varying from 0 to 100 μM in the presence and absence of 2.5 µM of ATV. After fluorescence quantification, the Michaelis-Menten constant (K_m_) and maximum velocity (V_max_) were calculated by non-linear regression using GraphPad Prism 9. The value was presented as mean ± standard deviation (SD).

### 2.4. Molecular docking procedure

The crystallographic structure of M^pro^ was obtained from Protein Data Bank (PDB), with access code 7K40 [18]. The chemical structure for ATV was built and minimized in terms of energy by Density Functional Theory (DFT) via Spartan’18 software (Wavefunction, Inc., Irvine, CA, USA) [19]. The molecular docking calculations were performed with GOLD 2020.2 software (Cambridge Crystallographic Data Center Software Ltd., CCDC, CB2 1EZ, UK) [20]. Hydrogen atoms were added to the protease following tautomeric states and ionization data, which are inferred by the GOLD 2020.2 software at pH 7.4. The number of genetic operations (crossing, migration, mutation) during the search procedure was set as 100,000. Redocking studies were carried out with the crystallographic ligand boceprevir (GC376, PDB code: 7K40), obtaining the lowest root mean square deviation (RMSD) value by ChemPLP function. It was defined 8 □ radius around the active binding site and the figures were generated with PyMOL Delano Scientific LLC software (DeLano Scientific LLC: San Carlos, CA, USA) [21].

### 2.5. Yield-reduction assays and virus titration

Calu-3 cells (2.0 × 10^5^ cells/well) were infected with multiplicity of infection (MOI) of 0.1 and 0.001 for SARS-CoV-2 B.1 lineage and MOI of 0.1 for SARS-CoV-2 gamma strain. Infection was performed in 96-well plates (Nalge Nunc Int, Rochester, New York, USA) for 1 h at 37°C in 5 % of CO_2_. Inoculum was removed and cells were incubated different concentrations of ATV (0.00, 0.60, 1.25, 2.50, 5.00, and 10.0 µM) or RDV (0.00, 0.0001, 0.001, 0.01, 0.10, 0.50, 1.0, 5.0, and 10.0, µM) in DMEM with 10% FBS. After 48 h, the supernatants were harvested, and infectious virus titers were quantified by plaque forming assays according to previous publications [7,22,23].

To perform the virus titration, Vero cells (2.0 × 10^4^ cell/well) in 96-well plates (Nalge Nunc Int, Rochester, New York, USA) were infected with log-based dilutions of the yield reduction assays’ supernatants for 1 h at 37°C in 5% of CO_2_. After the incubation, medium containing 1.8% CMC with 5% FBS was added and incubated at 37°C with 5% CO_2_ for 72 h. The cells were fixed with 10% formaldehyde in PBS and stained with a 0.04% solution of crystal violet in 70% methanol. The virus titers were calculated by scoring the plaque-forming unit – (PFU/mL) and a non-linear regression analysis of the dose-response curves was also performed to calculate the 50% effective concentration (EC_50_). All experiments were carried out at least three independent times, including a minimum of two technical replicates in each assay, and each data was analyzed from Prism GraphPad software 8.0 (Windows GraphPad Software, San Diego, California USA). The value was presented as mean ± standard deviation (SD).

### 2.6. Cytotoxic assays

Vero cells (2.0 × 10^4^ cell/well) were treated for 3 days with different concentrations of ATV or RDV (ranging from 1 to 600 μM) as previously described by us [7,23]. The 50% cytotoxic concentration (CC50) was calculated by a non-linear regression analysis from a dose–response curve. All experiments were carried out at least three independent times and each data was analyzed from Prism GraphPad software 8.0 (Windows GraphPad Software, San Diego, California USA). The results were presented as mean ± standard deviation (SD). The selectivity indexes (SI) for ATV and RDV were calculated through the ratio between CC_50_ and EC_50_ values.

### 2.7. Pharmacokinetic assays

ATV’s concentration in the plasma and lungs of adult Swiss-Webster mice (8-15 weeks) was evaluated over time. Animals were treated with an oral dose of 60 mg/kg of ATV for 12-time intervals - 00:05, 00:10, 00:20, 00:40, 01:00, 02:00, 03:00, 04:00, 06:00, 08:00, 10:00, and 12:00h. Each treatment group had 5 animals. After these periods of times, total blood and lungs were collected. Plasma was obtained by blood centrifugation at 8,000 x g for 15 minutes. Time zero was obtained by analyzing the matrix pool (blood plasma) of untreated animals. The pharmacokinetic parameters of half-life (t_1/2_, h), time until maximum concentration is reached (t_max_, h), maximum compound concentration (C_max_, ng/mL), and area under the curve (AUC, h.ng/mL) were determined. The experiments performed in this section were approved by the Committee on the Use of Laboratory Animals of the Oswaldo Cruz Foundation (CEUA-FIOCRUZ, license L003/21).

### 2.8. In vivo assays – Mice treatment and infections

Experiments with transgenic mice expressing human ACE-2 receptor (K18-hACE2-mice), were performed in Animal Biosafety Level 3 (ABSL-3) multiuser facility, according to the animal welfare guidelines of the Ethics Committee of Animal Experimentation (CEUA-INCa, License 005/2021) and WHO guidelines [14]. The animals were obtained from the Oswaldo Cruz Foundation breeding colony and maintained with free access to food and water at 29– 30°C under a controlled 12 h light/dark cycle. Experiments were performed during the light phase of the cycle.

For infection procedures, mice were anaesthetized with 60 mg/kg of ketamine and 4 mg/kg of xylazine and inoculated intranasally with DMEM high glucose (MOCK), or 10^5^ PFU of SARS-CoV-2 gamma strain in 10 μl of DMEM high glucose. It was used 6 mice per experimental group: MOCK (non-infected); SARS-CoV-2-infected without treatment (NIL) and SARS-CoV-2-infected and treated with ATV. The animals were treated with a daily dose of 60 mg/kg of ATV for seven days.

The animals were monitored daily during seven days for survival and body-weight analysis. In the case of weight loss higher than 20% euthanasia was performed to alleviate animal suffering. In the last day, the bronchoalveolar lavage (BAL) from both lungs was harvested by washing the lungs once with 1 mL of cold PBS. After centrifugation of BAL (1500 rpm for 5 minutes), the pellet was used for total and differential leukocytes counts (diluted in Turk’s 2% acetic acid fluid) using a Neubauer chamber. The lactate dehydrogenase (LDH) quantification was performed with centrifuged BAL supernatant to evaluated cell death (CytoTox96, Promega, USA). Differential cell counts were performed by cytospin (Cytospin3; centrifugation of 350xg for 5 minutes at room temperature) and stained by the May-Grünwald-Giemsa method.

After BAL harvesting, lungs were perfused with 20 mL of saline solution to remove the circulating blood. Lungs were then collected, pottered, and homogenized in 500 μL of a phosphatase and protease inhibitor cocktail Complete, mini EDTA-free Roche Applied Science (Mannheim, Germany) for 30 sec, using an Ultra-Turrax Disperser T-10 basic IKA (Guangzhou, China).

### 2.9. Quantification of viral RNA

The viral RNA from samples collected in the *in vivo* assays was quantified through quantitative reverse transcription polymerase chain reaction (RT-PCR). Total RNA was extracted using QIAamp Viral RNA (Qiagen), according to manufacturer’s instructions. Quantitative RT-PCR was performed using Quanti Tect Probe RT-PCR Kit (Qiagen) in a StepOne Plus™ Real-Time PCR System (Thermo Fisher Scientific). Amplifications were carried out in 15 μL reaction mixtures containing 2× reaction mix buffer, 50 μM of each primer, 10 μM of the probe, and 5 μL of RNA template. Primers, probes, and cycling conditions recommended by the Centers for Disease Control and Prevention (CDC) protocol were used to detect the SARS-CoV-2 (CDC 2020). Amplification of the housekeeping gene glyceraldehyde-3-phosphate dehydrogenase (GAPDH) was used as a reference for the number of cells. The cycle threshold (CT) values for this target were compared to those obtained with different cell quantities (10^7^ to 10^2^), for calibration.

### 2.10. Measurements of inflammatory mediators and cell death

The levels of IL-6, TNF-α, KC, and PF4 were quantified in BAL samples from uninfected (MOCK), infected without treatment (NIL), and infected and treated animals by ELISA, using specific kits and following the manufacturer’s instructions (R&D Systems). Cell death was determined according to the activity of LDH in the BAL as previously described in the section 2.7.

### 2.11. Histological procedure

Histological features related to the injury caused by SARS-CoV-2 infection were analyzed in the lungs of K18-hACE2 mice. Inflammatory and vascular infiltrates and evidence of cell degeneration was evaluated to characterize the level of the tissue damage. The collected material was fixed with formaldehyde (4%), dehydrated and embedded in paraffin to the obtention of tissue slices through the use of a microtome. The slices were fixed and stained with hematoxylin and eosin for microphotographs analysis.

### 2.12. Clotting time

Human blood samples were collected from healthy donors in 3.8% trisodium citrate (9:1, v/v), and platelet-poor plasma was obtained by centrifugation at 3,000 rpm for 10 min. Plasma (100 μL) was incubated with 1 μL of ATV at various concentrations (diluted in DMSO) for 2 min at 310K. Plasma clotting was initiated by the addition of 100 μL of 25 mM CaCl_2_, and the time for clot formation was recorded on a KC-4 Delta coagulometer (Tcoag, Ireland). Time for clot formation was recorded in triplicates.

## 3. Results

### 3.1. Enzymatic and cell-based assays for ATV in SARS-CoV-2 D614G and gamma strains

To advance on details on how ATV inhibits SARS-CoV-2 M^pro^, we initially performed a dose-dependent inhibition curve. ATV was slightly less potent than the positive control boceprevir (GC376) [24] (Figure 1A). The Morrison’s inhibitory constant (K_i_) values for GC376 and ATV were 208 ± 0.15 and 703 ± 79 nM, respectively. Next, we tested ATV against various concentrations of M^pro^ substrate. We observed that maximum velocity (V^max^) values in the presence and absence of ATV were not different, while the Michaelis-Menten constant (K^m^) values increased significantly in the presence of this drug (Figure 1B), indicating a competitive inhibition profile. Our results are specific to M^pro^ because, differently from the positive control GRL0617, ATV did not inhibit SARS-CoV-2 papain-like protease (PL^pro^) (Figure 1C). Since structural improvements on the M^pro^ active site were determined more recently [25,26], SARS-CoV-2 molecular docking calculations were carried out to test if a catalytic water (H_2_O_cat_) was required for ATV action. Since the highest docking score value was obtained in the presence of H_2_O_cat_, molecular docking calculation suggested a dependence of ATV potency of water content into the catalytic site of M^pro^ (Figure 1D).

**Figure 1.**
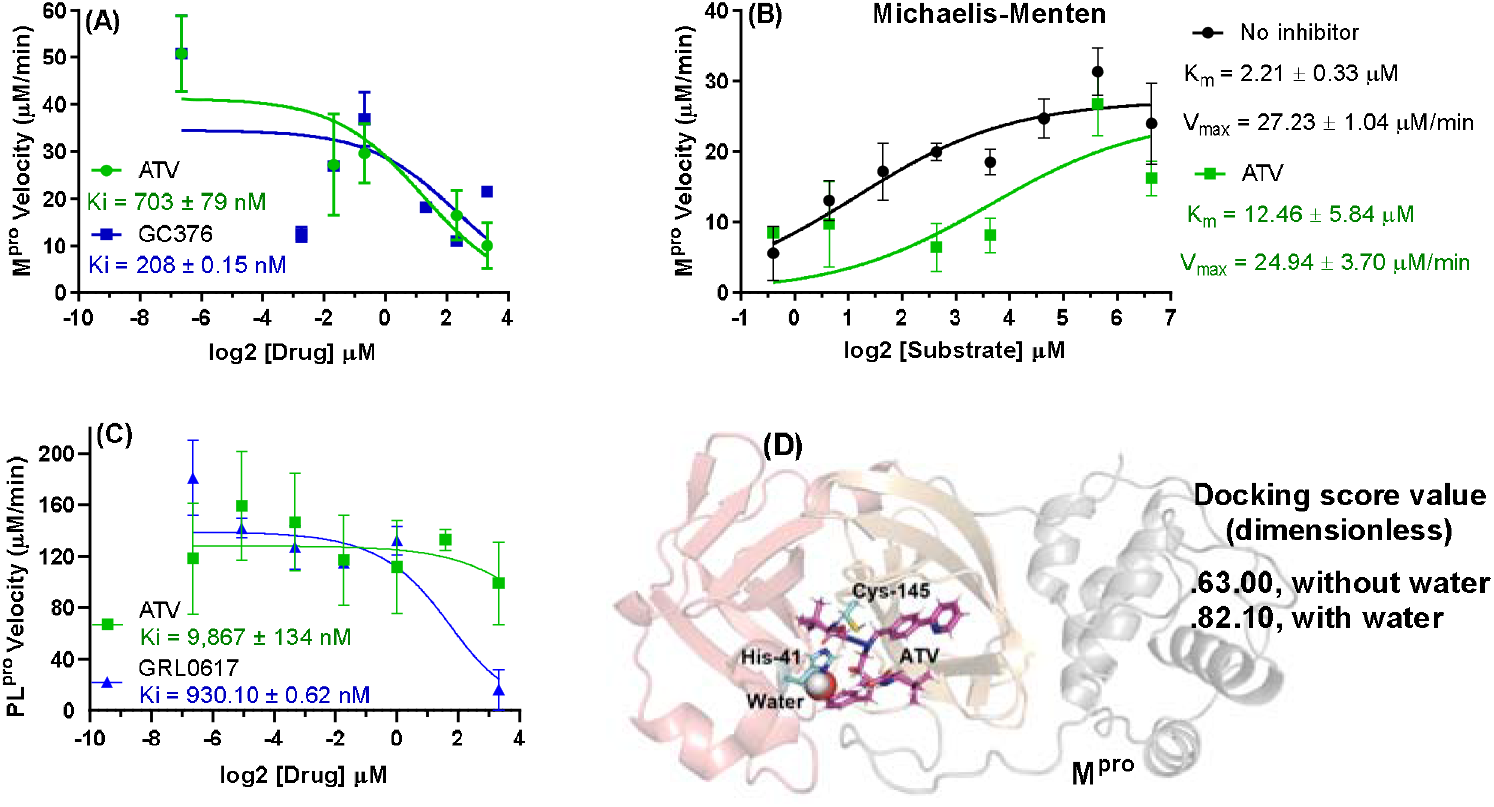
(A) The ATV and boceprevir (GC376, positive control) activity on 88.8 nM M^pro^ velocity at 0-10 μM of inhibitor (protease assay kit #79955-1, BPS Biosciences, USA). (B) Michaelis-Menten plot for 88.8 nM M^pro^ incubated overnight in assay buffer with substrate concentrations from 0 to 100 μM in the presence and absence of 2.5 µM of ATV. (C) The ATV and GRL-617 (positive control) activity on 100 nM PL^pro^ velocity at 0-10 µM of inhibitor (protease assay kit #79995-1, BPS Biosciences, USA). (D) The 3D representation of the best docking pose for ATV into M^pro^ catalytic site in the presence of the catalytic water (H_2_O_cat_) obtained with GOLD 2020.2 software (Cambridge Crystallographic Data Centre, Cambridge, CB2 1EZ, UK) using ChemPLP as scoring function. For better interpretation the M^pro^ structure was represented only in the monomeric form with the domains I, II, and III in light red, orange, and gray, respectively. The ATV and catalytic amino acid residues are in stick representation in pink and cyan, respectively, while H_2_O_cat_ is represented as sphere.

The competitive inhibition implies that M^pro^’s substrate concentration may affect its susceptibility to ATV. Thus, we tested if ATV’s potency is also affected in type II pneumocyte cell line (Calu-3) infected with different SARS-CoV-2 MOIs. In cell-based assays, the intracellular concentration of M^pro^’s substrate would be proportional to the virus input. Indeed, ATV’s EC_50_ value for the SARS-CoV-2 B.1 lineage varied in a MOI-dependent way (Table 1), similarly to remdesivir (RDV) (Table 1), a competitive inhibitor of the SARS-CoV-2 RNA polymerase complex under clinical use [27,28]. The SARS-CoV-2 gamma VoC is more susceptible to ATV and RDV than its predecessor strain B.1 (Table 1). Since the CC_50_ values of 312 ± 8 and 512 ± 30 µM for ATV and RDV, respectively, their selective index (SI) values were consistent with an adequate safety profile *in vitro* (Table 1).

**Table 1.**
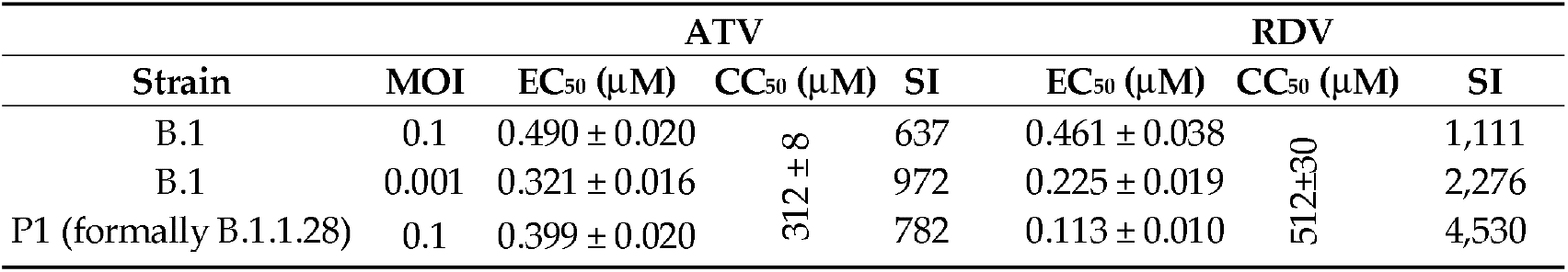
In vitro pharmacological parameters for ATV and RDV against SARS-CoV-2 variants in Calu-3 cells.

### 3.2. In vivo antiviral activity of ATV against SARS-CoV-2 gamma variant

We first aimed evaluate ATV’s pharmacokinetics profile over time in the plasma and lungs of Swiss-Webster mice treated with 60 mg/kg of this drug, a dosage equivalent to its plasma exposure in humans under treatment against HIV. Upon treatment with 60 mg/kg, ATV concentration in the plasma was similar to the standard treatment of 300 mg in humans (Figure 2A) [29]. Interestingly, ATV seems to be concentrated in the lung of the treated animals (Figure 2B).

**Figure 2.**
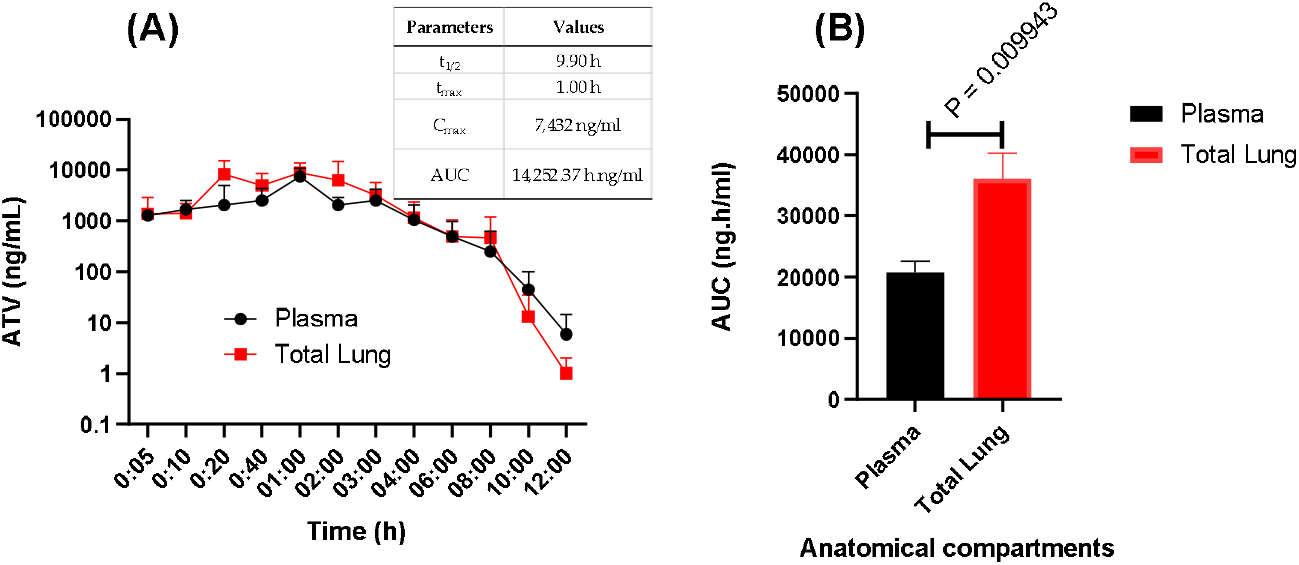
Pharmacokinetics of atazanavir (ATV) in mice. Swiss-Webster mice at 8-15 weeks of age were orally treated with 60 mg/kg of ATV. (A) At indicated time points, concentration of ATV was measured in the plasma and in the lungs. The insert in panel A represents the pharmacokinetic parameters in the plasma. (B) The area under the curve (AUC) for the anatomical compartments were registered.

Next, we infected K18-hACE2-transgenic mice with SARS-CoV-2 gamma VoC and treated them with a daily oral dose of 60 mg/kg ATV, initiating 12h after infection. Whereas the infection kills all animals within 6 days, a statistically significant increase in animal survival was observed in the infected and ATV-treated mice (Figure 3A). ATV protected the mice to continue to lose weight at the 6^th^ day after infection (Figures 3B). In the bronchoalveolar lavage (BAL) of the treated animals, ATV significantly decreased SARS-CoV-2 RNA levels (Figure 3C), cell death - based on lactate dehydrogenase (LDH) activity (Figure 3D), and cell-based inflammation – based on cells counts of polymorphonuclear and mononuclear cells (Figure 3E and F).

**Figure 3.**
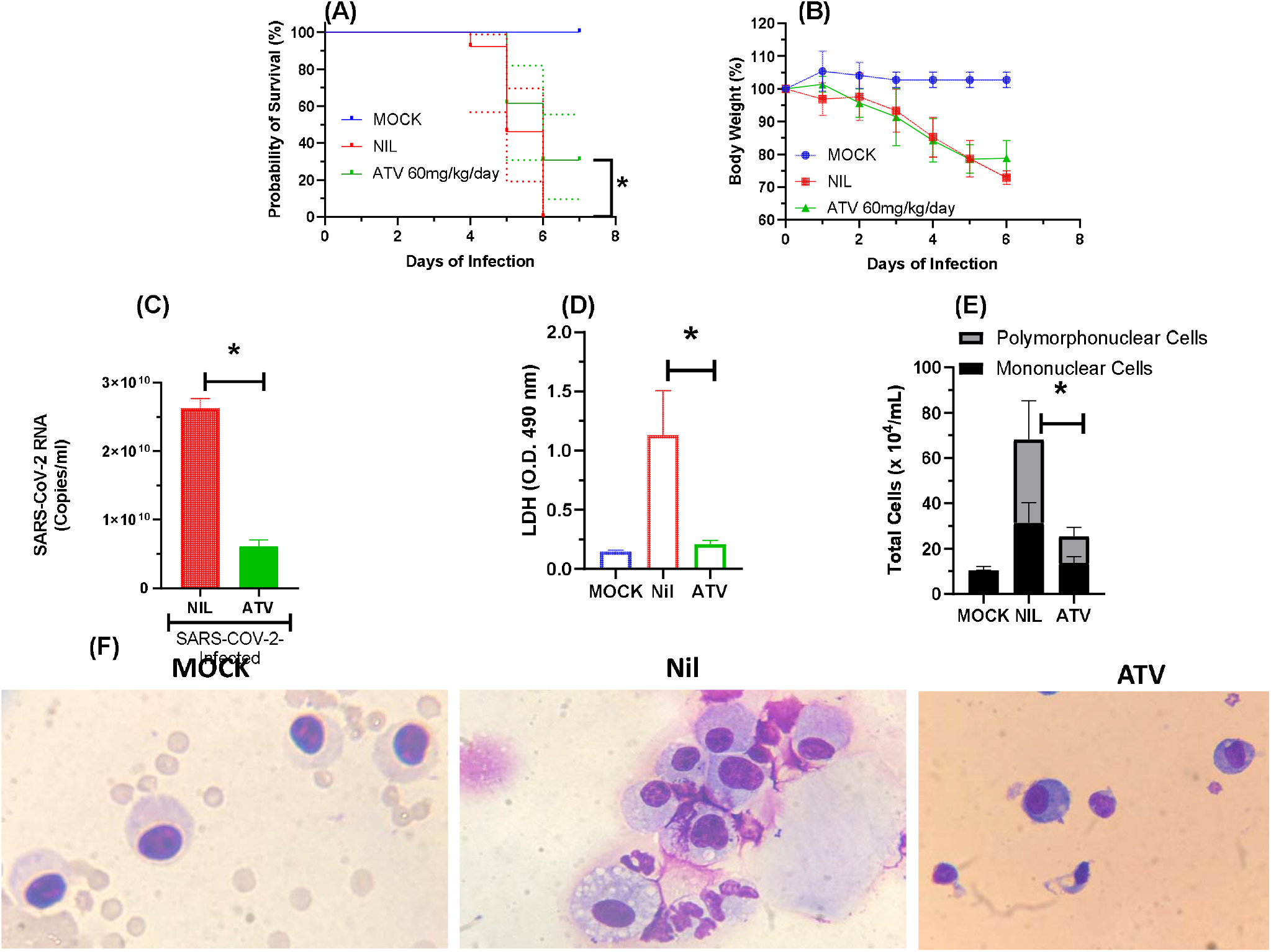
ATV protected K18-hACE2-transgenic mice infected with SARS-CoV-2 from mortality. The survival (A), and weight variation (B) of the different experimental groups: non-infected and non-treated (MOCK) or SARS-CoV-2-infected and non-treated (NIL) SARS-CoV-2-infected and treated with ATV. After 12h of infection the treated group received the first of a daily dose of 60 mg/kg of ATV. The animals were treated for six days. Survival was statistically assessed by Log-rank (Mentel-Cox) test, where * p < 0.05. The viral load (C), LDH levels (D), polymorphonuclear and mononuclear cells counts (E) and immunocytochemical staining (F) were assessed in the BAL six days after infection in the indicated experimental groups. All the analysis were performed with 6 mice/group.

ATV reduced the SARS-CoV-2-induced IL-6 levels in the BAL and in the lungs of treated animals (Figure 4). In the lung, levels of TNF-α and KC were also reduced in the infected/untreated over untreated mice (Figure 4). These results are in line with our previous description that ATV decreases the levels of SARS-CoV-2-induced pro-inflammatory cytokines in monocytes [7] and virus-triggered pyroptosis [30]. Moreover, SARS-CoV-2 infection provoked severe lung injury, leading to hemorrhage, and shrinking of the lobe, bronchiole, and alveoli (Figure 5), which was reduced by ATV. This protection is the consequence of the direct antiviral and anti-inflammatory activity of ATV, since this molecule could not prevent hemorrhage as an anti-clotting agent (Figure S1, Supplementary Material).

**Figure 4.**
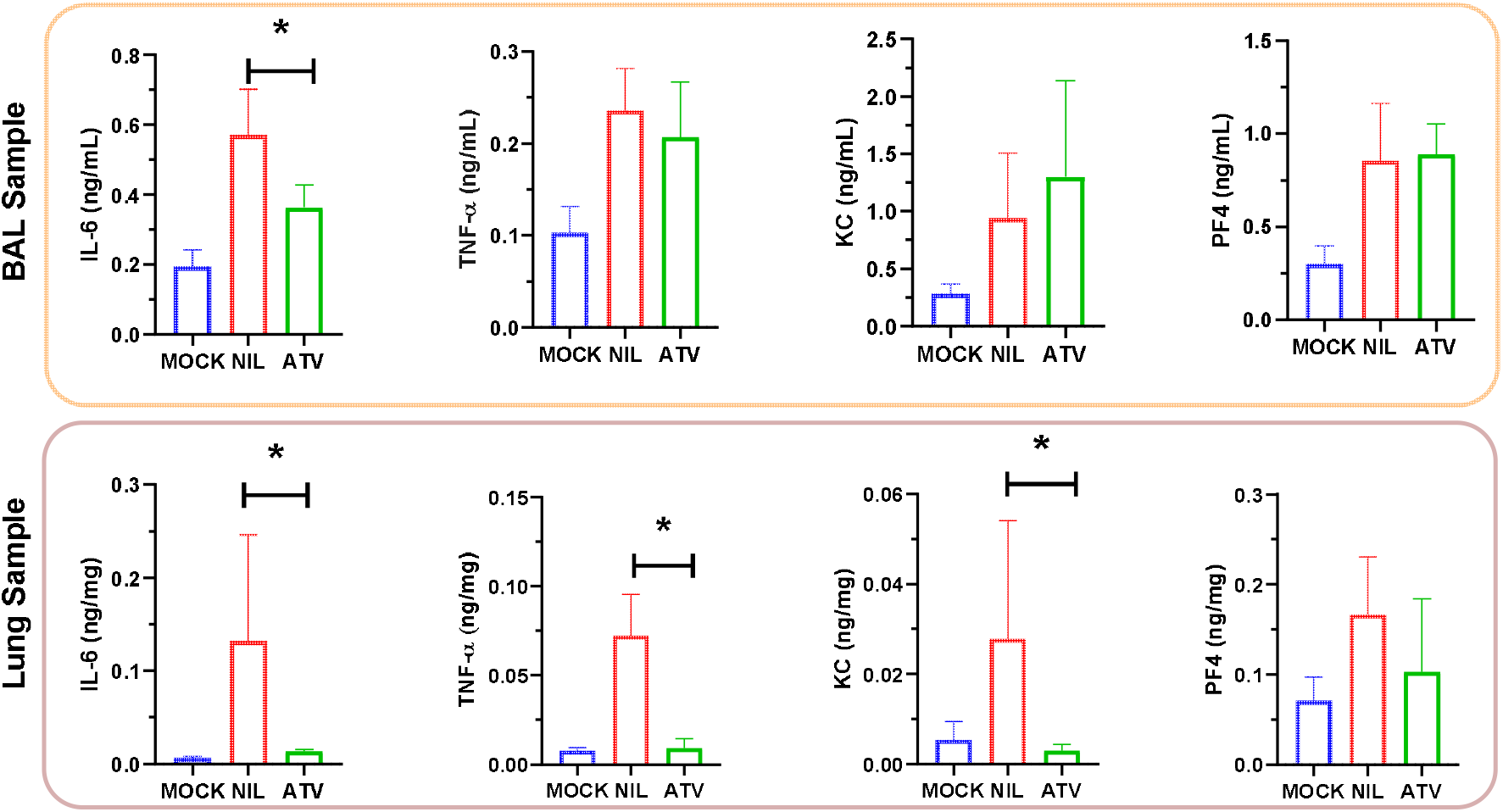
The proinflammatory content in terms of cytokines IL-6, TNF-α, KC, and PF4 in BAL and lung samples. * p < 0.05

**Figure 5.**
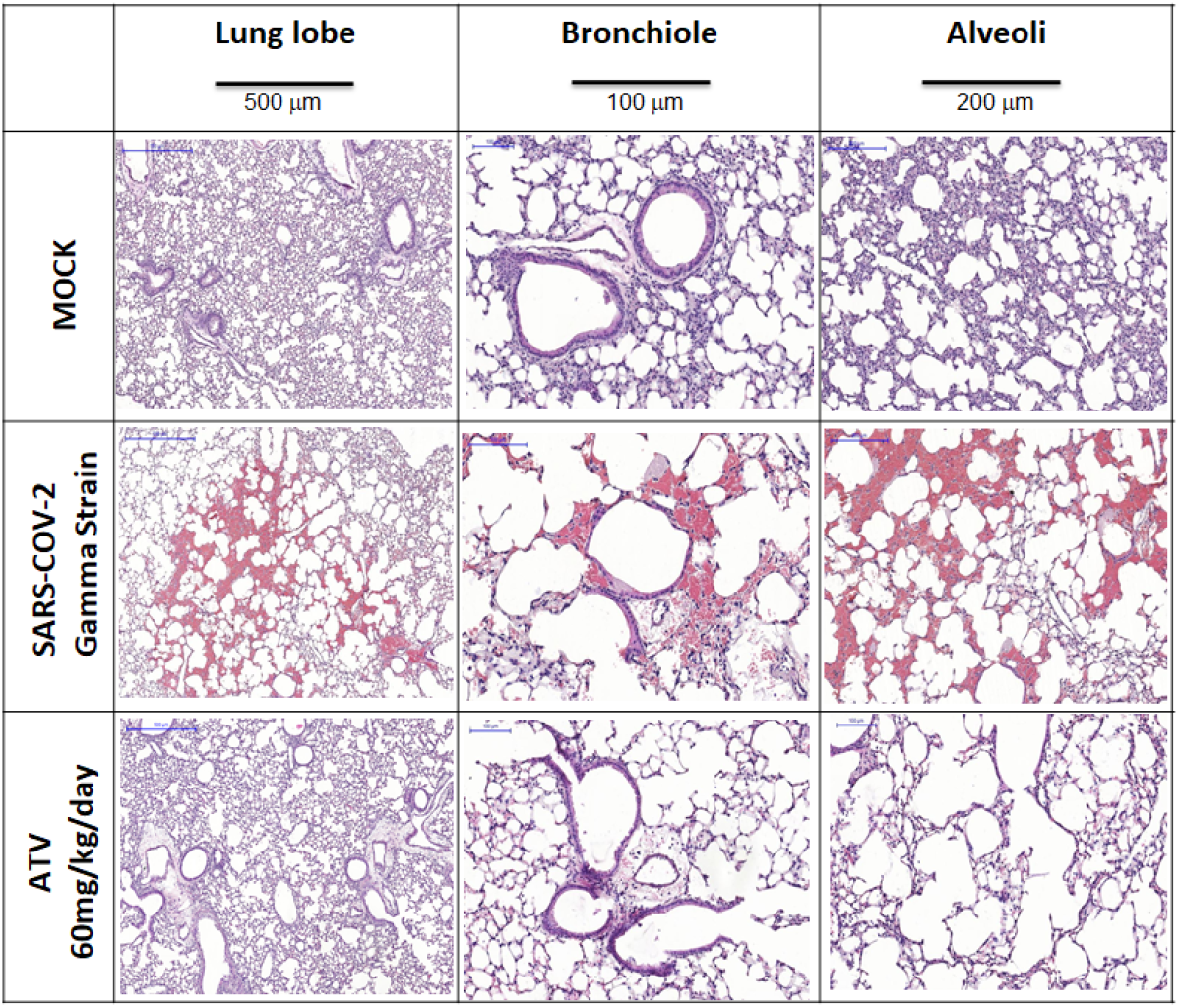
Microphotographs for histology of lung lobe, bronchiole, and alveoli samples from K18-hACE2-transgenic mice non-infected (MOCK) and infected with SARS-CoV-2 gamma strain without (NIL) and treated with ATV upon the six treated days.

## Discussion

Repurposing of clinically approved drugs was considered an accelerated strategy to combat SARS-CoV-2 infections [31,32]. However, limited clinical benefit has been documented for most repurposed drugs, whereas in the meantime orally available antiviral drugs against COVID-19 demonstrated clinical efficacy to reduce hospitalization, such as molnupiravir [33] and PAXLOVID™ [12]. Molnupiravir and PAXLOVID™ respectively target the viral RNA synthesis and M^pro^. Besides detailed mechanism of action, these compounds fulfilled pre-clinical steps of investigation, from cell-based to animal models, to allow further clinical development to be conducted at target plasmatic concentration. Here, we performed these steps for ATV. The comprehension that ATV inhibits competitively M^pro^ and requires a catalytic water may allow further development of anti-SARS-CoV-2 analogs, that act differently than PAXLOVID™, a covalent inhibitor of M^pro^ by interaction with the catalytic cysteine (Cys-145) residue [34].

We originally demonstrated that ATV inhibited SARS-CoV-2 M^pro^ preparations and virus replication [7]. The controversial effects of ATV on SARS-CoV-2 M^pro^ are documented in the literature [10,12]. In line with studies that demonstrated SARS-Cov-2 susceptibility to ATV, we demonstrated here that M^pro^ was inhibited by this drug, with K^i^ 1.5-fold higher than boceprevir (GC376) and in a competitive fashion, increasing the enzyme’s K_m_ more than 6-fold. It is unlikely that the other SARS-CoV-2 protease, PL^pro^, was targeted by ATV in these assays, because its K_i_ for PL^pro^ was above the threshold of *in vitro* inhibition.

The amino acid residue His-41 of M^pro^ requires water to catalyze proteolytic cleavage [25,26] and ATV targets this moiety. The presence of 20% glycerol in enzyme assay mixture reduces the water content and prevents ATV activity, that is why Ma and Wang [13,35] could not identify this HIV protease inhibitor as a potential antiviral against COVID-19 - whereas Li et al. [10] reached results similarly to ours. Our enzymatic assay includes bovine serum albumin (BSA) to avoid non-specific binding of small molecules to M^pro^. In our M^pro^ FRET-based enzymatic assay, reactions were allowed to occur overnight to increase the sensitivity, instead of 1h [13]. Although one might argue that a longer reaction time will increase the rate of false positive results, the following cell-based assays provide an additional layer of evidence that ATV activity against SARS-CoV-2 is credible.

Considering the substrate-dependent inhibitory effect of ATV, we hypothesized ATV’s potency in cell-based assays would be influenced by varying the virus input, because higher numbers of virus particles should translate into increased quantities of M^pro^ substrate. To test this hypothesis, Calu-3 cells, which is more closely resemble type II pneumocytes than A549 and Vero cells [36], were infected with a 100-fold different multiplicity of infections (MOI) of SARS-CoV-2, B.1 lineage, and treated with ATV or RDV, a competitive inhibitor of viral RNA synthesis. The MOI-dependent inhibition was consistent for ATV and RDV. The similarity in the cell-based potency of ATV to its K_i_ for M^pro^ further reinforces the conclusion that M^pro^ is the target of ATV *in vitro*.

The emerging Brazilian SARS-CoV-2 gamma variant, initially detected in the state of Amazonas, was responsible for a public health calamity, spreading rapidly in Brazil and considered as one of variant of concern by WHO. The mutations found in gamma variant have been associated with increased transmissibility, higher viral load, propensity for immune evasion, and SARS-CoV-2 reinfection [37,38]. ATV presented adequate inhibitory profile against the predecessor strain of SARS-CoV-2 and also the gamma variant *in vitro*. Based on that and considering the lethality of gamma variant on K-18 mice model, we next performed *in vivo* experiments.

At plasma exposures similar to humans [29], ATV enhanced by 30% the survival of K18-hACE2-transgenic mice infected with SARS-CoV-2 gamma strain and decreased virus-induced cell death and inflammation. Despite there is not a statical difference in the body weight between the untreated and treated groups, the protective effect of ATV was viewed in relation to the significantly decrease of SARS-CoV-2 RNA levels in the bronchoalveolar lavage (BAL), as well as a significant reduction in the number of mononuclear leukocytes, polymorphonuclear leukocytes and LDH levels. In the BAL infected and treated animals displayed lower levels of IL-6, TNF-α, KC and PF4, compared to untreated mice. The in vivo anti-inflammatory of ATV observed here is in line with the ability of this molecule to inhibit pyroptosis in human primary monocytes for both BAL and lung samples clearly showing that ATV protect both cell death and inflammation [39].

Our study shows that ATV inhibits SARS-COV-2 M^pro^ with a mechanism of action different than PAXLOVID™, showing ATV/M^pro^ interface could give insights for further drug development. The *in vitro* and *in vivo* results reconfirm, in different magnitudes, SARS-CoV-2 susceptibility to ATV. Further ongoing clinical trials will determine if standard dose, used in HIV-treatment can prevent severe COVID-19 or if it is necessary higher doses [40]. According to ATV’s monography [29], doses three times higher could be used for shorter periods of time, when compared to the life-lasting HIV treatment.

## Author Contributions

Conceptualization, T.M.L.S. and P.T.B.; methodology, O.A.C., C.Q.S., A.C.F., M.M., N.F.-R., J.R.T., and T.M.L.S.; software, O.A.C.; validation, O.A.C., T.M.L.S., R.Q.M., P.T.B., H.C.C.-F.-N., and J.P.B.V.; formal analysis, O.A.C., C.Q.S., A.C.F., M.M., N.F.-R., J.R.T., D.P.P., G.P.E.S., L.B.F., H.M.P., A.S.C., J.C.d’Ávila, L.V., and R.Q.M.; investigation, O.A.C., T.M.L.S., C.Q.S., A.C.F., M.M., N.F.-R., and J.R.T.; resources, T.M.L.S., J.P.B.V., P.T.B., and H.C.C.-F.-N.; data curation, O.A.C. and T.M.L.S.; writing—original draft preparation, O.A.C. and T.M.L.S.; writing—review and editing, O.A.C., T.M.L.S., C.Q.S., A.C.F., M.M., N.F.-R., J.R.T., R.Q.M., J.P.B.V., and H.C.C.-F.-N.; visualization, O.A.C. and T.M.L.S.; supervision, T.M.L.S., R.Q.M., J.P.B.V., P.T.B., and H.C.C.-F.-N.; project administration, T.M.L.S.; funding acquisition, T.M.L.S., J.P.B.V., P.T.B., and H.C.C.-F.-N. All authors have read and agreed to the published version of the manuscript.

## Institutional Review Board Statement

The *in vivo* studies were conducted according to the both guidelines of Committee on the Use of Laboratory Animals of the Oswaldo Cruz Foundation (CEUA-FIOCRUZ) with license L003/21 approved in 01/04/2020 and to the animal welfare guidelines of the Ethics Committee of Animal Experimentation from National Cancer Institute of Brazil (CEUA-INCA) with license 005/2021 approved in 03/05/2021.

## Informed Consent Statement

Not applicable.

## Acknowledgments

The authors acknowledgement the funding Brazilian agencies CNPq, Faperj, and CAPES and also Dra. Patricia Machado Rodrigues e Silva Martins and Dra Tatiana Paula Teixeira Ferreira from Fiocruz for the microphotographs analysis. O.A.C. also thanks Dr. Dumith Chequer Bou-Habib and *Fundação para o Desenvolvimento Científico e Tecnológico em Saúde* (FIOTEC) both from Oswaldo Cruz Foundation for the grant VPPCB-005-FIO-20.

## SUPPLEMENTARY MATERIAL

**Figure S1.**
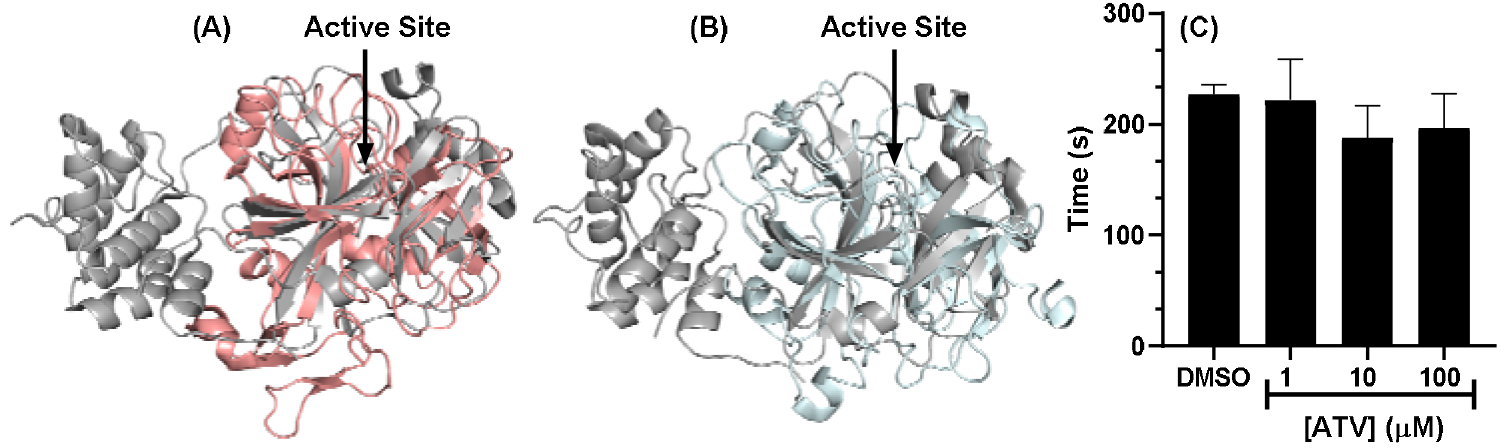
Superposition of the monomeric unit of M^pro^ (in gray, PDB code 7K40) with (A) FXa (in salmon, PDB code 2P16) and (B) thrombin (in cyan, PDB code 1KTS). For better interpretation the catalytic water (H_2_O_cat_) of M^pro^ is not shown. (C) Fibrin formation trial without and in the presence of three concentrations of ATV.

